# Understanding immune-driven brain aging by human brain organoid microphysiological analysis platform

**DOI:** 10.1101/2022.01.19.476989

**Authors:** Sunghwa Song, Zheng Ao, Hongwei Cai, Xiang Li, Yifei Miao, Zhuhao Wu, Jonathan Krzesniak, Mingxia Gu, Luke P. Lee, Feng Guo

**Affiliations:** Department of Intelligent Systems Engineering, Indiana University, Bloomington, IN 47405, United States; Perinatal Institute, Division of Pulmonary Biology, Cincinnati Children’s Hospital Medical Center, Cincinnati, OH 45229, United States; Center for Stem Cell and Organoid Medicine, CuSTOM, Division of Developmental Biology, Cincinnati Children’s Hospital Medical Center, Cincinnati, OH 45229, United States; Harvard Institute of Medicine, Harvard Medical School, Harvard University, Brigham and Women’s Hospital, Boston, MA, 02115, USA; Department of Bioengineering, Department of Electrical Engineering and Computer Science, University of California at Berkeley, Berkeley, CA, USA; Department of Biophysics, Institute of Quantum Biophysics, Sungkyunkwan University, Suwon, Korea

**Author notes:** Corresponding (F.G.) or (L.P.L.).

**Keywords:** Microfluidics, Brain Organoid, Aging, Neuro-Immune Interaction, Inflammaging

## Abstract

The aging of the immune system drives systemic aging and the pathogenesis of age-related diseases. However, a significant knowledge gap remains in understanding immune-driven aging, especially in brain aging, due to the limited current *in vitro* models of neuro-immune interaction. Here we report the development of a human brain organoid microphysiological analysis platform (MAP) to discover the dynamic process of immune-driven brain aging. We create the organoid MAP by 3D printing that can confine organoid growth and perfuse oxygen and nutrients (and immune cells) to generate standardized human cortical organoids that promote viability, maturation, and commitment to human forebrain identity. Dynamic rocking flow is incorporated for the platform that allows us to perfuse primary monocytes from young (20 to 30-year-old) and aged (>60-year-old) donors and culture human cortical organoids for modeling and analyzing the aged immune cell interacting organoid tissues systematically. We discovered the aged monocytes had increased infiltration and promoted the expression of aging-related markers (e.g., p16 in astrocytes neighboring to monocytes) within human cortical organoids, indicating that aged monocytes may drive brain aging. We believe that our human brain organoid MAP provides promising solutions for basic research and translational applications in aging, neuroimmunological diseases, autoimmune disorders, and cancers.

## Introduction

Aging is a complex process with the accumulation of unrepaired molecular and cellular damage.^1^ During aging, chronic, sterile, low-grade inflammation, named ‘inflammaging’, contributes to the pathogenesis of age-related diseases such as Alzheimer’s disease.^2, 3^ Aging of the immune system, or immunosenescence, drives systemic aging, including brain and other organisms.^4^ Taking one of the most important immune cell types in blood, bone marrow-derived monocytes can infiltrate the aged and/or inflamed brain and get activated to secrete inflammatory molecules (e.g., TNF-α), contributing to the inflammation in the aged brain.^5–7^ Tremendous interests have been attracted to studying immune-driven brain aging, e.g., aged monocytes impacting brain aging and aging-associated neural diseases, but little has been explored yet.^8–10^

Several models have been employed to investigate neuro-immune interaction for understanding inflammation and age-related diseases. 2D in vitro cultures are simple, high throughput, and widely used to model the interaction of neuronal and immune cells. In contrast, the 2D cultures lack the complex brain tissue architectures and neuro-immune microenvironments of an in vivo brain. Animals have been demonstrated as an excellent model to study neuroimmunology at different physiological and pathological conditions. However, the animal model is costly, time-consuming, and has difficulty fully capturing human biology due to their genomic and epigenomic differences from humans.^11, 12^ Superior to 2D cultures and animals, human brain organoids, 3D in vitro brain-like tissues derived from human stem cells, can recapitulate some critical features of the human brain, including cellular identities, brain structures, and microenvironments, and neural activities, bringing new insights to study neuro-immune interaction in aging and age-related diseases.^13–15^ For example, human brain organoids have been developed to model the microglia-mediated neuroinflammation for understanding the etiology of Alzheimer’s disease, substance use disorder, and other diseases.^16–18^ However, current organoids suffer from poor penetration of oxygen/nutrients, variable reproducibility, and lack of immune components. Thus, there is an emergent and unmet need for the reproducible generation of standardized organoids with immune components to model neuro-immune interaction.

Recently, microfluidics and microfabrication technologies have been used for improving the organoid cultures and 3D in vitro cultures from aspects including standardization, throughput, function, etc.^19–28^ For example, microfluidic droplets have been used to encapsulate cells into water-in-oil droplets or microgels for high throughput generation of uniformed human brain organoids and tumor spheroids.^29–31^ Based on our development of acoustofluidics,^32–36^ our group has adapted this technology to assemble cells for the massive and scaffold-free generation of standardized 3D cell cultures and assembloids.^37–40^ The microfabricated perfusion devices have been developed to perfuse oxygen, nutrients, and/or chemicals into organoids or 3D in vitro cultures to improve the generation, development, and maturation/vascularization of kidney organoids and brain organoids.^41–44^ Pioneering efforts have also been made to develop automated microfluidic systems and mini-bioreactors to generate and track many organoids and 3D cultures (e.g., gastrointestinal, tumor, and brain organoids) for high throughput screening applications.^45–49^ Despite current microfluidic devices and systems having significantly improved the contemporary culture of organoids, there is still lacking the user-friendly and straightforward platforms that can reproducibly generate standardized organoids with immune components and allow the time-lapse imaging of neuro-immune interaction for studying aging and age-related diseases.

Herein, we developed innovative organoid microphysiological analysis to generate a standardized organoid per device and enable their interaction with immune cells. Our MAP has several advantages: (1) Our device design allows us to confine a developing organoid into the pancake shape as well as perfuse oxygen nutrients for avoiding hypoxia and necrosis; (2) Our method can mimic blood vessels to flow immune cells (e.g., monocytes) toward organoids for studying immune-organoid interaction; (3) In contrast to the traditional organoid culture methods (well-plates on a rocking machine, or bulk bioreactors), our technology will immobilize one organoid per device to avoid the organoid fusion as well as improve the standardization of organoid cultures. (4) Our 3D printed devices are simple, low-cost, user-friendly, and compatible with commonly used well-plates and lab equipment. They can be easily adapted with the current organoid protocol for high throughout the investigation of immune-organoid interaction. We used our devices to model immune-driven brain aging using primary human monocytes as a proof-of-concept. Interestingly, our results indicate aged monocytes may induce aging phenotype inside human brain organoids (high expression p16 of astrocytes within organoid) without external genetic manipulation.

## Results

### An organoid microphysiological analysis platform to model immune-organoid interaction

To model the immune-driven brain aging (**Figure 1a**), we developed an organoid microphysiological analysis platform (MAP) to investigate human brain organoid interaction with macrophages from old and young donors in a standardized and user-friendly manner. Our MAP device (design details in **Fig.S1**) consists of two components (**Figure 1b, c**): (1) a hollow and meshed tubular perfusable scaffold connected with two medium reservoirs for supporting the organoid growth along the tubular scaffold surface, flowing in oxygen and nutrients, and mimicking blood vessels to flow immune cells (e.g., monocytes) toward organoids, and (2) an organoid holder on a coverslip for holding the organoids within well-plates. A developing organoid can grow into a pancake-shape structure within the confined space between the tubular perfusable scaffold and the organoid holder (within 400 μm). Once combined with rocking flows, our MAP can avoid the necrosis and/or hypoxia of the on-chip cultured organoids and facility the noninvasive introduction of immune cells and enable the time-lapse imaging of dynamic immune-organoid interaction. Our MAP may be widely used to culture different types of organoids once incorporated with specific organoid developmental protocols.

**Figure 1.**
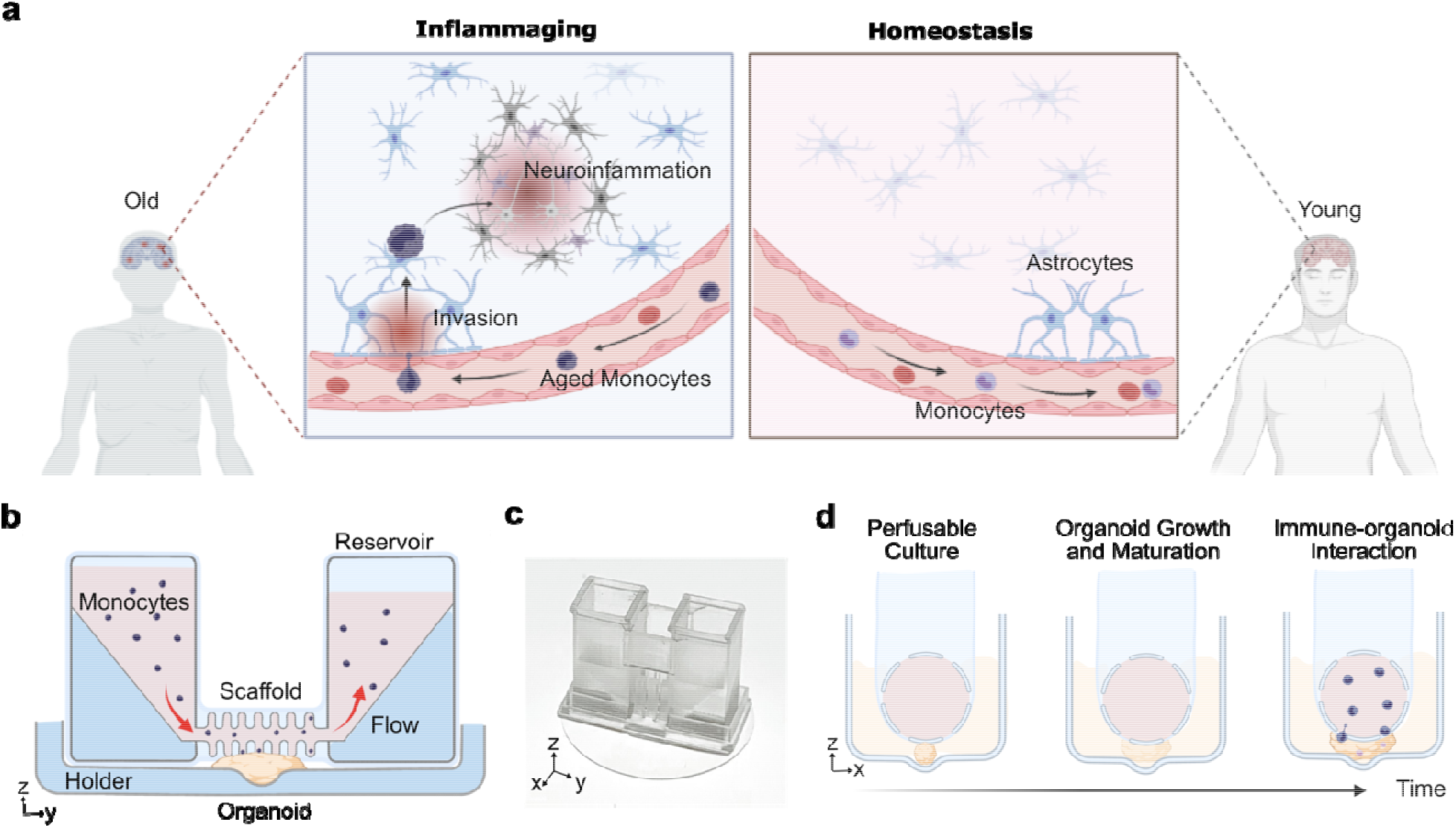
An organoid microphysiological analysis platform to model immune-driven brain aging. **(a)** Schematics show the 3D printed devices for modeling immune-driven brain aging by culturing and analyzing the interaction of primary monocytes (from young and aged donors) and human brain organoids. **(b)** Images of 3D printed devices within a well-plate and **a** single device with an organoid holder and a tubular perfusable scaffold connected with two medium reservoirs.

### On-chip culture of human cortical organoids under flows

Before applying our MAP for the organoid culture experiment, we generated and optimized the flow within our MAP using simulation and experimental approaches. Our simulation results described the 3D distribution (**Figure 2a**, left) and the side view (**Figure 2a**, right) of flow profiles within the MAP device (rocking angle = 15, and rocking frequency = 0.1 rpm, Supplementary discussion 1). With the theoretical prediction, we further measured the dependence of the flow speed on the rocking angle (dots in **Figure 2b**, with rocking frequency = 0.1 rpm), which matches our simulation result (a red dashed line in **Figure 2b**). We adopted a culture protocol to generate human cortical organoids (hCOs) on our devices (**Figure 2c**). Briefly, embryonic bodies (EBs) with a diameter of about 300 μm were assembled from human embryonic stem cells (WA01, Wicell) with SB-431542 and XAV-939 to inhibit TGF and Wnt/β-catenin signaling for neural induction and patterning. The EBs were then transferred to the organoid holder of the devices within a wellplate at day 9, embedded in Matrigel *in situ*, and cultured inside the device for 15 days till forming a hollow structure surrounding the mesh tubular scaffold (**Figure 2d**). Various growth factors, such as BDNF, EGF, or FGF, were added to the medium for enhancing the differentiation, maturation, and survival of neurons. On day 24, the devices within the well-plate were finally cultured on a rocking platform to generate inner lumen fluid flow. Meanwhile, ascorbic acid and cAMP were added to support the differentiation of neural progenitor cells into mature neurons and support organoid growth.

**Figure 2.**
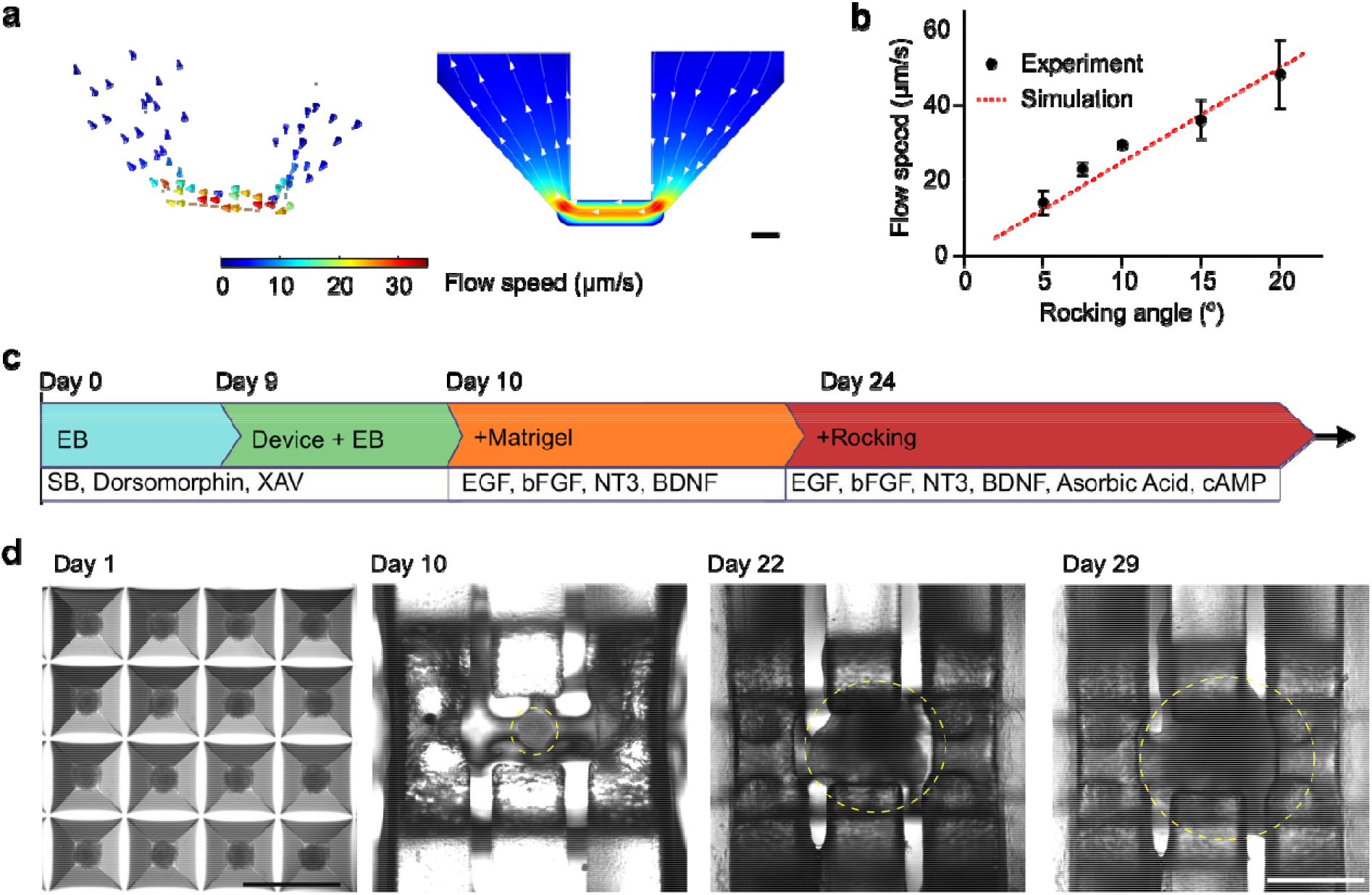
On-chip culture of human cortical organoids under flows. **(a)** The simulation results indicate the 3D distribution (left) and side view (right) of flow profiles within the device. **(b)** The experimental results describe the flow speed under different rocking angles (rocking frequency = 0.1 rpm). **(c)** The illustration shows the protocol and time of the on-chip culture of human cortical brain organoids. **(d)** Top-down view images of organoids during the differentiation and on-chip culture process (organoids in yellow dashed circles). Scale bar: 1 mm

### Characterization of on-chip cultured organoids

We characterized the development of on-chip cultured hCOs using viability tests, immunostaining, and sequencing approaches. We first used live/dead staining to test the viability of on-chip cultured hCOs on days 9, 16, and 24, respectively. As expected, with the support of medium perfusion, on-chip cultured hCOs maintained very high viabilities during a prolonged culture time (**Figure 3a**). We then confirmed the proper development and differentiation of on-chip cultured hCOs using PAX6 (neural progenitor) and MAP2 (neuron) staining. We found that the on-chip cultured hCOs successfully developed with proper ventricular/subventricular zones (VZ/SVZ) indicated by PAX6 staining as well as cortical plates indicated by MAP2+ neurons surrounding VZ/SVZ. (**Figure 3b**). We further compared gene expression of on-chip cultured hCOs with conventional hCOs (cultured on orbital shaker) using bulk RNA sequencing. Three on-chip cultured hCOs and three conventional hCOs were used to determine the change of gene expression related to neural development. To visualize the differentially expressed genes in our on-chip cultured and conventional hCOs, we first performed a hierarchical clustering analysis of the gene expression profiles. We found that the gene expression profiles of these two different groups clustered with each other (**Figure 3c**), indicating that our devices broadly impact the phenotypes of developing hCOs. Upon detailed examination of differentially expressed genes, we found that genes related to forebrain development (GSX2, FOXG1, and NKK2-1)^50^, deep layer cortical neuron development (BCL11B)^51^, and GABAergic neuron development (GAD1 and GABRA1)^52^ were significantly enriched in the on-chip cultured hCOs than conventional hCOs. Meanwhile, non-forebrain fate markers such as RAX and PMCH,^50, 53^ frequently found in the hypothalamus, were lower expressed in on-chip cultured hCOs than conventional hCOs (**Figure 3d**). These results indicated that the on-chip culture could facilitate forebrain fate commitment in hCOs, which is consistent with our previous report.^54^ We then further performed Kyoto Encyclopedia of Genes and Genomes (KEGG) pathway enrichment analysis to identify significant pathways impacted by on-chip culture. We found that the differentially expressed genes were significantly enriched in diverse synaptic development pathways such as GABAergic, dopaminergic, cholinergic, and glutamatergic synapses pathways (**Figure 3e**). Overall, these results show that our devices could facilitate forebrain fate commitment, neuron and synapse maturation in hCOs.

**Figure 3.**
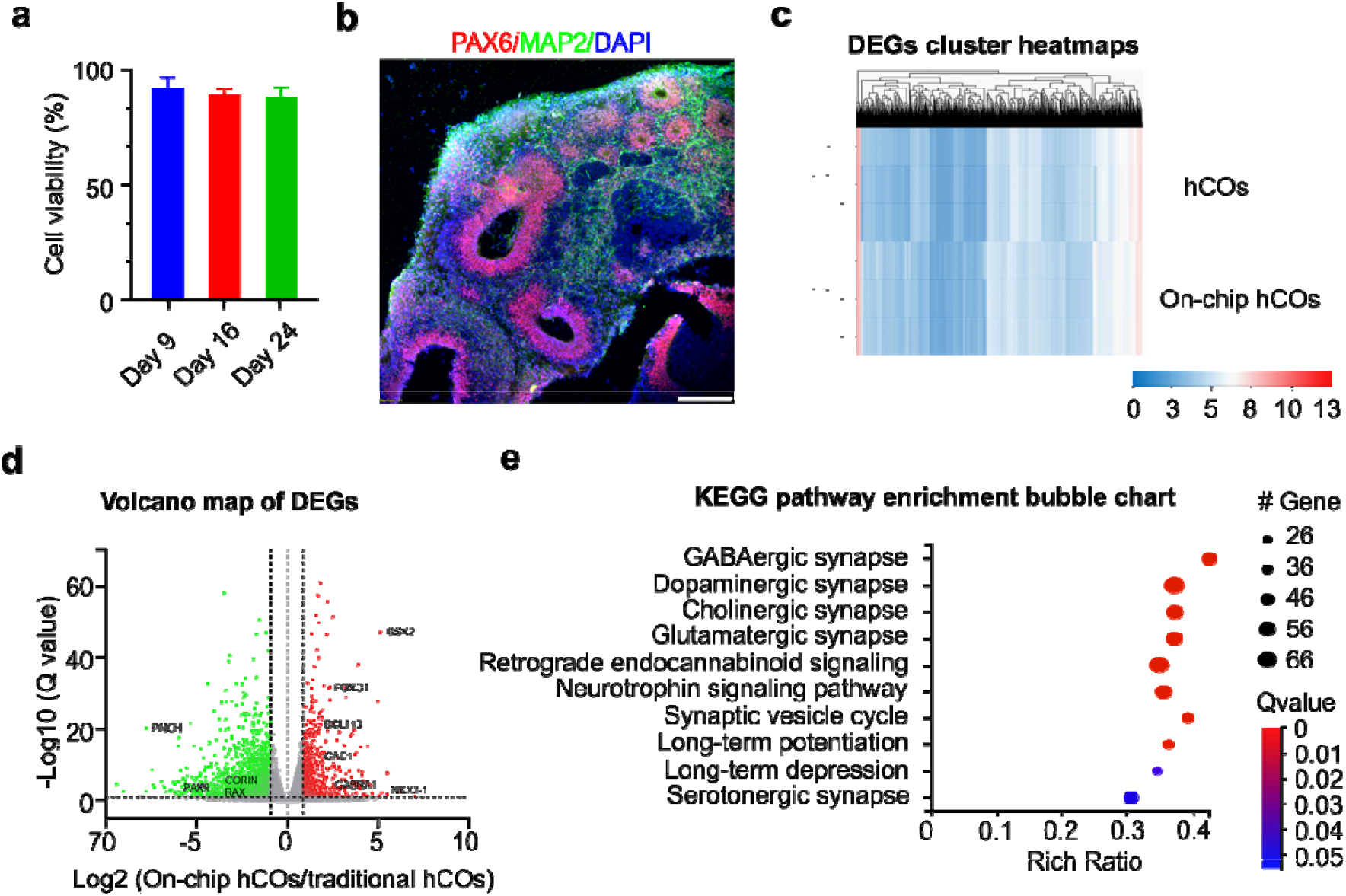
Characterization of on-chip cultured organoids. **(a)** Viability of on-chip cultured human cortical organoids (on-chip hCOs) during long-term culture. **(b)** Cross-section staining showing VZ/SVZ (PAX6 and MAP2 neuron) distribution inside the on-chip hCO at day 29. **(c)** Cluster heatmaps of differentially expressed genes (DEGs) in the on-chip hCOs and the conventional human cortical organoids (hCOs). **(d)** Volcano map of DEGs with red (increased expression) and green (decreased expression) dots. **(e)** KEGG pathway enrichment bubble chart showing high-level functions and utilities in the on-chip hCOs than the conventional hCOs. Scale bar: 500 μm.

### On-chip investigation of monocyte infiltrating organoid

In the aged brain, monocytes may have a greater capacity to infiltrate the brain on the brain-blood barrier (BBB) during injury or neurological disorders.^55^ Particularly, animal experiments showed that aged mice harbor expanded CCR2+ monocytes post-injury/chronic inflammation that can infiltrate the brain based on MCP1/CCR2 chemotaxis signaling.^56^ Post infiltration, monocytes may induce pro-inflammatory responses in neighboring glial cells such as astrocytes. To investigate the differential infiltration capacity of young and aged monocytes into hCOs, we perfused the primary monocytes isolated from PBMCs into the on-chip cultured hCO (**Movie S1**). We collected the old monocytes (oMs) from the PBMCs from the aged (>60 years old) donors and obtained young monocytes (yMs) from the young (20-30 years old) donors. After perfusing oMs and yMs into devices separately, we recorded the monocyte infiltrating organoid on-chip (**Figure 4a**). We found the infiltrated cell number of oMs was about 3 times compared to that of yMs at 24 hours post perfusion (**Figure 4b**), indicating oMs have significantly higher infiltration capacity than yMs. Upon further qRT-PCR analysis, we discovered that the on-chip cultured oMs and hCO (oMs+hCO) expressed higher levels of monocyte chemoattractant genes such as MCP1 and MCP3 than the on-chip cultured yMs and hCO (yMs+hCO) (**Figure 4c**). This indicated that the increased infiltration capacity of oMs could also be attributed to MCP1/CCR2 signaling axis.

**Figure 4.**
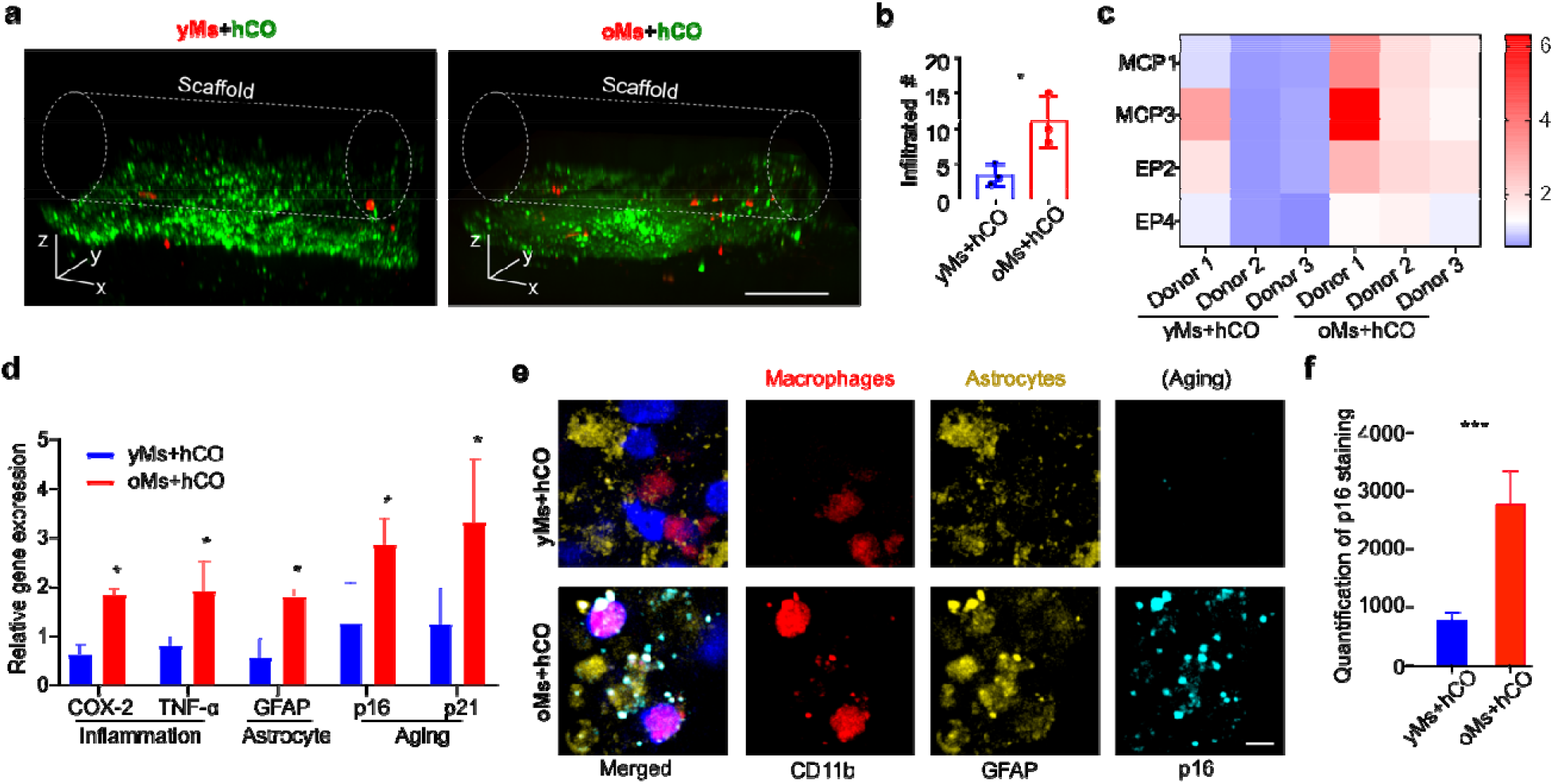
Characterization of monocyte-driven brain organoid aging. **(a)** Comparison of infiltration capacity of monocytes isolated from old donors (> 60-year-old) into on-chip cultured human cortical organoid (oMs+hCO) with monocytes isolated from young donors (20 to 30-year-old) into on-chip cultured human cortical organoid (yMs+hCO). **(b)** Quantification of infiltrated monocytes (n=3, p<0.05) in ‘yMs+hCO’ and ‘oMs+hCO’ groups. Scale bar: 200 μm. **(c)** The plot of gene expression change (2^−ΔΔCT^) heatmap of the ‘yMs+hCO’ and ‘oMs+hCO’ groups on day 29 under the same experimental conditions. **(d)** Comparison of pro-inflammation genes (COX-2 and TNF-alpha) and senescence genes (p16^INK4a^ and p21^CIP1^) in on-chip cultured monocytes isolated from old donors (> 60-year-old) and human cortical organoid (oMs+hCO) and these in on-chip cultured monocytes isolated from young donors (20 to 30-year-old) and human cortical organoid (yMs+hCO) on day 29. 2^−ΔΔCT^ calculated as ‘delta Ct’ (ΔΔCt) of GAPDH and target gene normalized against on-chip cultured human cortical organoid only on the same day. **(e)** Immunofluorescence staining of the p16 expression in neighboring astrocytes monocytes in the ‘yMs+hCO’ and ‘oMs+hCO’ groups. Scale bar: 5 μm. **(f)** Quantification of p16 protein expression in GFAP+ astrocytes neighboring infiltrated monocytes using mean fluorescence intensity, p<0.005, n=30.

### On-chip investigation of monocyte-driven organoid aging

Cellular senescence is one of the hallmarks of aging, as indicated by senescence markers such as p16. Expression of p16 is driven by various genetic and environmental factors such as inflammation. Thus, we sought to investigate the effect of monocyte infiltration on hCO. Notably, the differential changes that young and old monocytes bring to hCO, regarding inflammation and aging markers. By qRT-PCR analysis of on-chip co-cultured monocytes and hCO, we discovered an upregulation of prostaglandin E2 (PGE2), COX-2, and TNF-alpha (**Figure 4d**), which are age-associated macrophage related proinflammation genes.^57, 58^, as well as upregulation of GFAP, as an indication of astrocyte activation within the oMs+hCO group.^59, 60^ Interestingly, we also found a significantly higher expression of aging-related markers including p16 and p21 in the oMs+hCO group. To further confirm that the increased p16 expression does not solely come from infiltrated oMs alone, we additionally performed immunofluorescence staining of p16 proteins. We found that p16 expression was higher in the astrocytes (GFAP+) neighboring infiltrated oMs (**Figure 4e**). Using mean fluorescence intensity, we quantified the p16 protein expression and found the p16 expression in astrocytes neighboring infiltrated monocytes within the oMs+hCO group was 2 times higher than that within the yMs+hCO group (**Figure 4f**). These results indicated a potential mechanism for infiltrated monocytes to induce p16 expression inside neighboring astrocytes to promote brain aging. Although a detailed mechanism must be elucidated, our findings shed light on a potential approach to inducing aging phenotype inside hCOs without external genetic manipulation.

## Discussion

We represent an organoid microphysiological analysis platform to model neuro-immune interaction in aging. By integrating organoid scaffold and rocking flow, our MAP allows the generation of standardized human cortical brain organoids without hypoxia and necrosis. The on-chip culture organoids showed significant forebrain fate commitment compared to the traditional culture. Additionally, due to the noninvasive incorporation of immune cells into organoids with unique tubular scaffold and rocking flow, our MAP supports the investigation of primary immune-organoid interaction on-chip, recapitulating the key features and processes of neuro-immune interaction in the aged brain. Our organoid chips device is simple, scalable, costeffective, and compatible with well-plates and lab settings. We, therefore, believe that our technology provides avenues to study immune-driven brain aging and restoring brain homeostasis in aging, with the coal of improving neurodegenerative functions.

## Methods

### Design and fabrication of 3D printed devices

The MAP devices were designed using AutoCAD software and fabricated using a 3D printer (Form 3B, Formlabs). The detailed design of the MAP devices was described at **Fig.S1**.

### Simulation and experimental measurement of flow profiles

The device’s flow profiles were simulated using Comsol Multiphysics (COMSOL group, details in **Discussion S1** and **Fig.S2**). Briefly, the rocking flow was affected using the ‘laminar flow’ physics module, and the liquid-scaffold boundary was considered non-slip. To model the fluid movement, ‘moving mesh’ was applied to the fluid domain, and the top air-liquid interface was set as a ‘free surface’ model. Half of the model was simulated with symmetric boundary conditions, and quantification results were obtained using the 2D model to save the computational energy. The experimental measurement of the flow speed was conducted by tracking and calculating dye transferring with the device under five different rocking angles (details in **Discussion S2** and **Fig.S3**). Briefly, the device was placed on the rocker at a neutral position. 30 μL of culture medium was added to both sides of the medium reservoir, with one side containing 0.83 mg/mL of Rhodamine B. The flow induced by the rocking cycle was videotaped at 30 frames per second with a fixed positioned camera secured on the rocker with a tripod. Video frames were analyzed with ImageJ. The flow speed was calculated by timing the moving dye front passing through a predetermined distance (3 mm) within the scaffold device.

### Culture of WA01 cells and THP-1 cells

The human embryonic stem cell line (WA01, WiCell) was maintained with mTeSR plus medium (Stemcell Technologies) on 6 well plates coated with Matrigel (Corning). The WA01 cells were passaged every 5-7 days with a medium change every other day and generated organoids under passage 42. The THP-1 (ATCC^®^ TIB-202™) cells, peripheral blood-derived monocytes, were cultured in RPMI 1640 (Gibco) supplemented with 10% fetal bovine serum (Gibco), 0.05 mM 2-mercaptoethanol (Sigma), and 100 U/mL Penicillin/Streptomycin (Gibco). Cells were maintained at 37 °C, 5% CO_2_ and passaged before the cell concentration reached 1 × 10^6^ cells/mL.

### Differentiation protocol of human cortical organoids

Human embryonic stem cells were dissociated into single cells using ReLeSR (Stemcell Technologies) treatment. Single cells were plated to ultra-low attachment 96 well plate (Corning) with AggreWell™ EB Formation Medium (Stemcell Technologies) with 10 μM Y-27632 (SelleckChem) (9,000 cells/well). After 24 hours, the medium was substituted by cortical organoid medium (1X DMEM/F12; 15% KOSR; 1X GlutaMax; 1X MEM-NEAA; 1X N2 supplement; 1X N2 supplement; 1X 2-Mercaptoethanol; 1X Penn/Strep) supplemented with 1μM SB-431542, 2μM Dorsomorphin and 2μM XAV939 for 9 days. Next, the aggregates were embedded in Matrigel and cultured in the medium (0.5X DMEM/F12; 0.5X Neuralbasal; 1X N2; 1X MEM-NEAA; 1X GlutaMax; 2.5μg/mL Insulin; 1X b-mercaptoethanol; 1X Penn/Strep; 1X 10ng/mL EGF, 10ng/mL FGF-2, 10ng/mL NT3 and 10ng/mL BDNF) supplemented with 1X B27 without Vitamin A for 14 days. Then the aggregates were maintained in the medium (supplemented with 1X B27 with Vitamin A, 0.2mM Ascorbic Acid, 0.2mM cAMP, and 1% Matrigel) with a medium change every other day. The composition details and medium change timelines are provided in **Table S1**.

### Cell viability test

To measure long-term cell viability, hCOs were stained with a LIVE/DEAD Viability/Cytotoxicity Kit (Invitrogen). Samples were incubated with a carboxyfluorescein succinimidyl ester (CFSE) and ethidium homodimer-1 (EthD-1) mixture for 30 minutes at 37 °C. After washing the samples twice with the fresh medium, the fluorescence microscopy of the two different dyes was visualized by an Olympus IX83 inverted motorized microscope.

### Bulk cell sequencing

RNA was extracted from hCOs using Qiagen RNeasy plus mini kit. The RNA samples were sent to BGI Americas for RNA-sequencing analysis. The RNA-seq data were analyzed and visualized using Dr. Tom’s data visualization software.

### Isolation of monocytes from human PBMCs

To use primary monocytes, human Peripheral Blood Mononuclear Cells (PBMCs) were isolated from human buffy coats of six different healthy young (n=3, <35) and elderly (n=3, >65) donors. The human buffy coats were first spun sown at 300 g for 10 minutes. To remove RBCs, the supernatant was discarded, and 5 ml of ACK lysing buffer was added to the pellet, which was shacked for 5 minutes. After adding 20 ml of HBSS, the tube was centrifuged at 300 g for 10 minutes. Several times of RBCs removal process were repeated until RBCs were removed enough. Monocytes were isolated from PBMCs using Classical Monocyte Isolation Kit according to manufacturer’s protocol (Miltenyi Biotec GmbH) with MS Column (Miltenyi Biotec GmbH). Typically, 5 x 10^4^ monocytes per milliliter of whole blood were obtained.

### On-chip coculture of monocytes and organoids

Th hCOs of day 9 were carefully collected and washed in the pre-warmed COM I with a cupped tip. After aspiration of the surrounding medium, organoids were carefully transferred to the center chamber with 6 μl of the Matrigel and incubated in a 37°C incubator for 30 minutes. The scaffold was inserted onto the center chamber with an additional fresh medium followed by incubation. The device was then maintained in the COM II until day 24 with a medium change every other day. When changed to COM III, the organoids on-chip was cultured onto a platform rocker (Infinity rocker, Next Advance) with 1 cycle/min in the incubator. To infiltrate monocytes into the device, a total of 1 x 10^5^ cells of monocytes were suspended in 20 ul of COM III and added on the on-chip cultured hCOs. After 2 hours of incubation, the upper component of the device with an additional medium was inserted on the hCOs.

### Immunofluorescence staining

Immunofluorescence staining was performed on sectioned samples directly on slides to characterize on-chip cultured hCOs. The samples were washed twice with 1XPBS to remove the O.C.T. After antigen retrieval for 15 minutes with 3N hydrochloric acid (HCl) treatment, the sections were rinsed twice with 1X PBS and then treated with 0.3% Triton-X100/5% normal goat serum in PBS for 1 hour. Next, primary antibodies were added to the slides in a diluted antibody solution and incubated overnight at 4°C within a dark, humidified slide box. The samples were rinsed 3 times with 1XPBS and incubated with secondary antibodies at RT for 1 hour, then the slides were washed 3 times with FBS. Coverslips were mounted with ProLong Gold Antifade Mountant with DAPI (Invitrogen). Detailed information on primary and secondary antibodies can be found in **Table S2**.

### Real-time qPCR analysis

The Gene expression profile of hCOs and on-chip cultured hCOs was evaluated by RT-qPCR. In brief, organoids were first washed twice with PBS and lysed using RNeasy Plus Mini Kit (Qiagen) for RNA extraction. RNA was reverse transcribed to ds-cDNA using the qScript cDNA synthesis kit (Quanta Biosciences). Applied Biosystems™ Power SYBR™ Green PCR Master Mix (Thermo Fisher) was used to carry out the real-time qPCR. Detailed information on primers can be found in **Table S3**. ΔΔCt method was performed to calculate relative expression (−ΔΔ CT), the delta Ct value of target gene between organoid and organoid chips, both normalized against Ct value of GAPDH. Each qPCR reaction was tested in triplicates, and 3 organoids/brain-in-a-chips were analyzed for each group. The means of Ct values are represented. Samples as “not detected” were denoted as a Ct value of 40.

### Statistical analysis

The statistic comparison of each group was performed via t-test with GraphPad Prism 7. The statistical significance of differences in values is denoted: *p<0.05, **p<0.01, ***p<0.005. ****p<0.001.

## Acknowledgment

S.S. and Z.A. contributed equally to this work. F.G. acknowledges Indiana University departmental start-up funds, and the National Institute of Health Awards (R03EB030331 and DP2AI160242). The authors also thank the Indiana University Imaging Center (NIH1S10OD024988-01) for using their instruments.

## References

1. T. B. L. Kirkwood, Cell, 2005, 120, 437–447.

2. C. Franceschi, M. Bonafè, S. Valensin, F. Olivieri, M. De Luca, E. Ottaviani and G. De Benedictis, Ann N Y Acad Sci, 2000, 908, 244–254.

3. P. L. Minciullo, A. Catalano, G. Mandraffino, M. Casciaro, A. Crucitti, G. Maltese, N. Morabito, A. Lasco, S. Gangemi and G. Basile, Arch Immunol Ther Ex, 2016, 64, 111–126.

4. M. J. Yousefzadeh, R. R. Flores, Y. Zhu, Z. C. Schmiechen, R. W. Brooks, C. E. Trussoni, Y. X. Cui, L. Angelini, K. A. Lee, S. J. McGowan, A. L. Burrack, D. Wang, Q. Dong, A. P. Lu, T. Sano, R. D. O’Kelly, C. A. McGuckian, J. I. Kato, M. P. Bank, E. A. Wade, S. P. S. Pillai, J. Klug, W. C. Ladiges, C. E. Burd, S. E. Lewis, N. F. LaRusso, N. V. Vo, Y. S. Wang, E. E. Kelley, J. Huard, I. M. Stromnes, P. D. Robbins and L. J. Niedernhofer, Nature, 2021, DOI: 10.1038/s41586-021-03547-7.

5. G. Naert and S. Rivest, J Mol Cell Biol, 2013, 5, 284–293.

6. F. Ginhoux and S. Jung, Nature Reviews Immunology, 2014, 14, 392–404.

7. S. Li, E. Y. Hayden, V. J. Garcia, D. T. Fuchs, J. Sheyn, D. A. Daley, A. Rentsendorj, T. Torbati, K. L. Black, U. Rutishauser, D. B. Teplow, Y. Koronyo and M. Koronyo-Hamaoui, Front Immunol, 2020, 11, 49.

8. C. L. Hsieh, E. C. Niemi, S. H. Wang, C. C. Lee, D. Bingham, J. S. Zhang, M. L. Cozen, I. Charo, E. J. Huang, J. L. Liu and M. C. Nakamura, Journal of Neurotrauma, 2014, 31, 1677–1688.

9. J. M. Morganti, T. D. Jopson, S. Liu, L. K. Riparip, C. K. Guandique, N. Gupta, A. R. Ferguson and S. Rosi, Journal of Neuroscience, 2015, 35, 748–760.

10. B. D. Semple, N. Bye, M. Rancan, J. M. Ziebell and M. C. Morganti-Kossmann, Journal of Cerebral Blood Flow and Metabolism, 2010, 30, 769–782.

11. P. E. C. Leite, M. R. Pereira, G. Harris, D. Pamies, L. M. G. dos Santos, J. M. Granjeiro, H. T. Hogberg, T. Hartung and L. Smirnova, Part Fibre Toxicol, 2019, 16.

12. K. Grenier, J. Kao and P. Diamandis, Mol Psychiatr, 2020, 25, 254–274.

13. D. Pamies, T. Hartung and H. T. Hogberg, Exp Biol Med, 2014, 239, 1096–1107.

14. M. Jurga, A. W. Lipkowski, B. Lukomska, L. Buzanska, K. Kurzepa, T. Sobanski, A. Habich, S. Coecke, B. Gajkowska and K. Domanska-Janik, Tissue Eng Part C-Me, 2009, 15, 365–372.

15. H. Y. Tan, H. Cho and L. P. Lee, Nat Biomed Eng, 2021, 5, 11–25.

16. P. R. Ormel, R. V. de Sa, E. J. van Bodegraven, H. Karst, O. Harschnitz, M. A. M. Sneeboer, L. E. Johansen, R. E. van Dijk, N. Scheefhals, A. B. van Berlekom, E. R. Martinez, S. Kling, H. D. MacGillavry, L. H. van den Berg, R. S. Kahn, E. M. Hol, L. D. de Witte and R. J. Pasterkamp, Nat Commun, 2018, 9.

17. Z. Ao, H. Cai, Z. Wu, S. Song, H. Karahan, B. Kim, H. C. Lu, J. Kim, K. Mackie and F. Guo, Lab Chip, 2021, DOI: 10.1039/d1lc00030f.

18. J. Penney, W. T. Ralvenius and L. H. Tsai, Mol Psychiatr, 2020, 25, 148–167.

19. S. E. Park, A. Georgescu and D. Huh, Science, 2019, 364, 960–965.

20. T. Takebe and J. M. Wells, Science, 2019, 364, 956–959.

21. A. Sontheimer-Phelps, B. A. Hassell and D. E. Ingber, Nature Reviews Cancer, 2019, 19, 65–81.

22. F. Yu, W. Hunziker and D. Choudhury, Micromachines-Basel, 2019, 10.

23. V. Velasco, S. A. Shariati and R. Esfandyarpour, Microsyst Nanoeng, 2020, 6.

24. Y. K. Shou, F. Liang, S. L. Xu and X. K. Li, Frontiers in Cell and Developmental Biology, 2020, 8.

25. F. Duzagac, G. Saorin, L. Memeo, V. Canzonieri and F. Rizzolio, Cancers, 2021, 13.

26. I. Khan, A. Prabhakar, C. Delepine, H. Tsang, V. Pham and M. Sur, Biomicrofluidics, 2021, 15.

27. Y. Q. Wang, L. Wang, Y. J. Zhu and J. H. Qin, Lab on a Chip, 2018, 18, 851–860.

28. B. Zhang, M. Montgomery, M. D. Chamberlain, S. Ogawa, A. Korolj, A. Pahnke, L. A. Wells, S. Masse, J. Kim, L. Reis, A. Momen, S. S. Nunes, A. R. Wheeler, K. Nanthakumar, G. Keller, M. V. Sefton and M. Radisic, Nat Mater, 2016, 15, 669–678.

29. H. Liu, Y. Wang, H. Wang, M. Zhao, T. Tao, X. Zhang and J. Qin, Advanced Science, 2020, 7, 1903739.

30. Z. Wu, Z. Gong, Z. Ao, J. Xu, H. Cai, M. Muhsen, S. Heaps, M. Bondesson, S.-S. Guo and F. Guo, ACS Applied Bio Materials, 2020, 3, 6273–6283.

31. H. Wang, H. T. Liu, X. Zhang, Y. Q. Wang, M. Q. Zhao, W. W. Chen and J. H. Qin, Acs Appl Mater Inter, 2021, 13, 3199–3208.

32. F. Guo, Y. Xie, S. Li, J. Lata, L. Ren, Z. Mao, B. Ren, M. Wu, A. Ozcelik and T. J. Huang, Lab on a Chip, 2015, 15, 4517–4523.

33. F. Guo, P. Li, J. B. French, Z. Mao, H. Zhao, S. Li, N. Nama, J. R. Fick, S. J. Benkovic and T. J. Huang, Proceedings of the National Academy of Sciences of the United States of America, 2015, 112, 43–48.

34. F. Guo, Z. Mao, Y. Chen, Z. Xie, J. P. Lata, P. Li, L. Ren, J. Liu, J. Yang, M. Dao, S. Suresh and T. J. Huang, Proc Natl Acad Sci U S A, 2016, 113, 1522–1527.

35. X. Ding, P. Li, S.-C. S. Lin, Z. S. Stratton, N. Nama, F. Guo, D. Slotcavage, X. Mao, J. Shi and F. Costanzo, Lab on a Chip, 2013, 13, 3626–3649.

36. A. Ozcelik, J. Rufo, F. Guo, Y. Gu, P. Li, J. Lata and T. J. Huang, Nat Methods, 2018, 15, 1021–1028.

37. Z. Ao, H. Cai, Z. Wu, J. Johnathon, H. Wang, K. Mackie and F. J. b. Guo, Lab Chip, 2020, 21, 688–699.

38. H. Cai, Z. Ao, L. Hu, Y. Moon, Z. Wu, H.-C. Lu, J. Kim and F. Guo, Analyst, 2020, 145, 6243–6253.

39. H. Cai, Z. Wu, Z. Ao, A. Nunez, B. Chen, L. Jiang, M. Bondesson and F. Guo, Biofabrication 2020, 12, 035025.

40. B. Chen, Y. Wu, Z. Ao, H. W. Cai, A. Nunez, Y. H. Liu, J. Foley, K. Nephew, X. B. Lu and F. Guo, Lab on a Chip, 2019, 19, 1755–1763.

41. K. A. Homan, N. Gupta, K. T. Kroll, D. B. Kolesky, M. Skylar-Scott, T. Miyoshi, D. Mau, M. T. Valerius, T. Ferrante, J. V. Bonventre, J. A. Lewis and R. Morizane, Nat Methods, 2019, 16, 255–+.

42. Y. Wang, L. Wang, Y. Zhu and J. Qin, Lab Chip, 2018, 18, 851–860.

43. M. Zhang, P. Wang, R. Luo, Y. Wang, Z. Li, Y. Guo, Y. Yao, M. Li, T. Tao and W. Chen, Advanced Science, 2021, 8, 2002928.

44. M. Nikolaev, O. Mitrofanova, N. Broguiere, S. Geraldo, D. Dutta, Y. Tabata, B. Elci, N. Brandenberg, I. Kolotuev, N. Gjorevski, H. Clevers and M. P. Lutolf, Nature, 2020, 585, 574–578.

45. B. Schuster, M. Junkin, S. S. Kashaf, I. Romero-Calvo, K. Kirby, J. Matthews, C. R. Weber, A. Rzhetsky, K. P. White and S. Tay, Nat Commun, 2020, 11.

46. J. A. Brassard, M. Nikolaev, T. Hübscher, M. Hofer and M. P. Lutolf, Nature Materials, 2021, 20, 22–29.

47. N. Brandenberg, S. Hoehnel, F. Kuttler, K. Homicsko, C. Ceroni, T. Ringel, N. Gjorevski, G. Schwank, G. Coukos, G. Turcatti and M. P. Lutolf, Nature Biomedical Engineering, 2020, 4, 863–874.

48. X. Y. Qian, H. N. Nguyen, M. M. Song, C. Hadiono, S. C. Ogden, C. Hammack, B. Yao, G. R. Hamersky, F. Jacob, C. Zhong, K. J. Yoon, W. Jeang, L. Lin, Y. J. Li, J. Thakor, D. A. Berg, C. Zhang, E. Kang, M. Chickering, D. Nauen, C. Y. Ho, Z. X. Wen, K. M. Christian, P. Y. Shi, B. J. Maher, H. Wu, P. Jin, H. L. Tang, H. J. Song and G. L. Ming, Cell, 2016, 165, 1238–1254.

49. H. Cai, Z. Ao, Z. Wu, S. Song, K. Mackie and F. Guo, Lab Chip, 2021, 21, 2194–2205.

50. S. J. Yoon, L. S. Elahi, A. M. Pasca, R. M. Marton, A. Gordon, O. Revah, Y. Miura, E. M. Walczak, G. M. Holdgate, H. C. Fan, J. R. Huguenard, D. H. Geschwind and S. P. Pasca, Nat Methods, 2019, 16, 75–78.

51. P. Arlotta, B. J. Molyneaux, J. Chen, J. Inoue, R. Kominami and J. D. Macklis, Neuron, 2005, 45, 207–221.

52. E. Gatta, A. Guidotti, V. Saudagar, D. R. Grayson, D. Aspesi, S. C. Pandey and G. Pinna, Int J Neuropsychoph, 2021, 24, 130–141.

53. S. Yoo, J. Kim, P. Lyu, T. V. Hoang, A. Ma, V. Trinh, W. N. Dai, L. Z. Jiang, P. Leavey, L. Duncan, J. K. Won, S. H. Park, J. Qian, S. P. Brown and S. Blackshaw, Science Advances, 2021, 7.

54. Z. Ao, H. Cai, Z. Wu, S. Song, H. Karahan, B. Kim, H.-C. Lu, J. Kim, K. Mackie and F. Guo, Lab on a Chip, 2021.

55. A. Chou, K. Krukowski, J. M. Morganti, L.-K. Riparip and S. Rosi, International journal of molecular sciences, 2018, 19, 1616.

56. J. M. Morganti, L.-K. Riparip, A. Chou, S. Liu, N. Gupta and S. Rosi, J Neuroinflammation, 2016, 13, 1–12.

57. E. S. Chambers, M. Vukmanovic-Stejic, B. B. Shih, H. Trahair, P. Subramanian, O. P. Devine, J. Glanville, D. Gilroy, M. H. Rustin and T. C. Freeman, Nature Aging, 2021, 1, 101–113.

58. S. Kyrkanides, A. H. Moore, J. A. Olschowka, J. C. Daeschner, J. P. Williams, J. T. Hansen and M. K. O’Banion, Molecular Brain Research, 2002, 104, 159–169.

59. S. A. Villeda, J. Luo, K. I. Mosher, B. Zou, M. Britschgi, G. Bieri, T. M. Stan, N. Fainberg, Z. Ding and A. Eggel, Nature, 2011, 477, 90–94.

