## Supplemental information for "Understanding immune-driven brain aging by human brain organoid microphysiological analysis platform"

**Contents:**

**Supplementary Figure**

- **Figure S1:** Device design of organoid microphysiological analysis platform.

**Supplementary Discussion**

- **Discussion S1:** Simulation of flow distribution on-chip.
- **Discussion S2:** Characterization of flow rate on-chip.

**Supplementary Tables**

- **Table S1:** Cortical organoid protocol medium compositions.
- **Table S2:** Antibody used in immunofluorescence staining.
- **Table S3:** Primer sequences for qPCR analysis.

**References**

**Supplementary Figure**

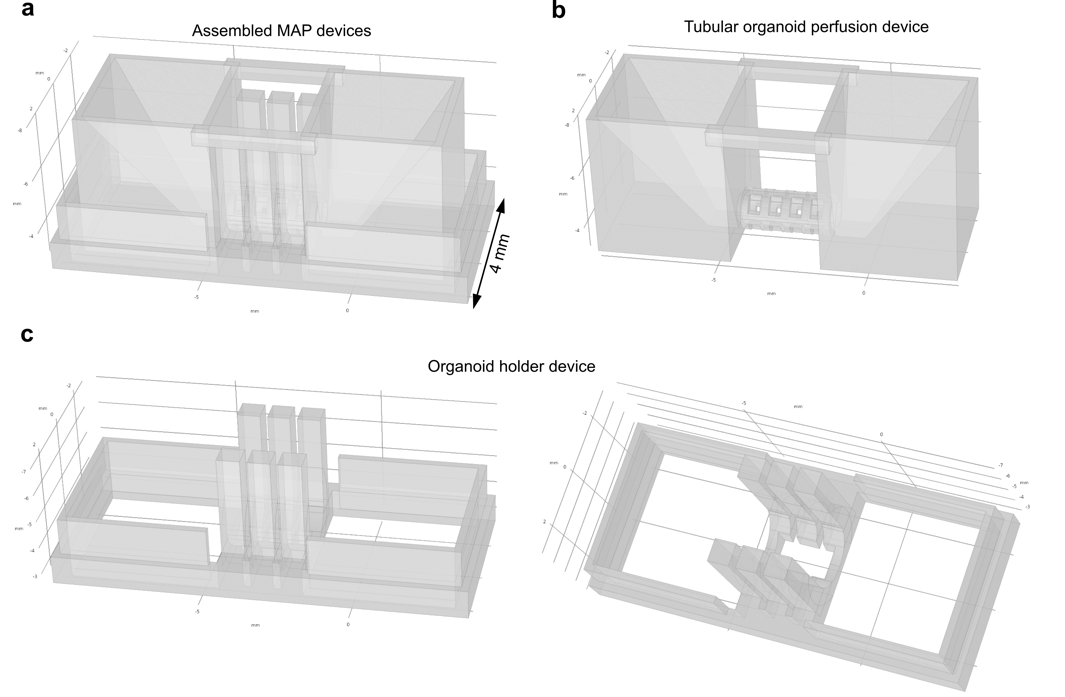

**Fig.S1. Device design of organoid microphysiological analysis platform.** (a) The design of assembled MAP devices with a tubular organoid perfusion device (upper) and an organoid holder device (bottom). (b) The design of tubular organoid perfusion device with a hollow and meshed tubular perfusable scaffold connected with two medium reservoirs. (c) The design of an organoid holder device on a coverslip.

**Supplementary Discussion**

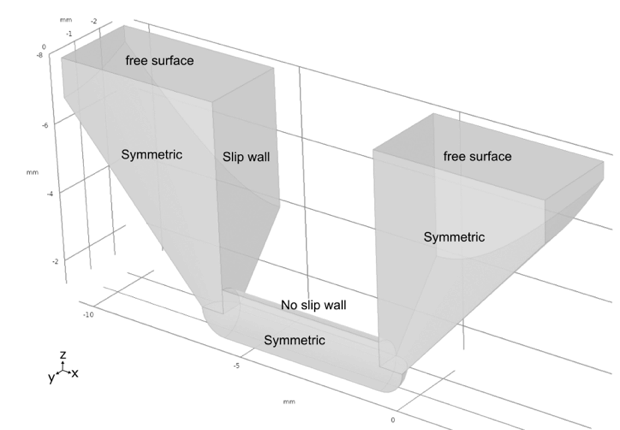
**Discussion S1: Simulation of flow distribution on-chip.** Simulation of rocking flow in the tubular device was conducted using Comsol Multiphysics (COMSOL group). To simplify the numerical simulation, only the liquid domain was considered in the simulation, while the 3D printed scaffold was consider by adding boundary condition as below **Fig.S2**.

**Fig.S2. Simulation domain and boundary conditions.** Half liquid domain with symmetric boundary condition is simulated. The air-liquid interface is set as “free surface” condition to model the liquid movement induced by rocking. The scaffold is considered as “slip wall”, except for the tubular part is “no slip wall”.

Briefly, the rocking liquid was simulated using ‘laminar flow’ physics module. The liquid-scaffold boundary of two side reservoir was considered as “slip wall” condition, while the tube part was considered as “no slip”. To model the liquid movement, ‘moving mesh’ was applied to the liquid domain and the top air-liquid interface was set as ‘free surface’ model. To save the computational energy, half part of the model was simulated with symmetric boundary condition and quantification results were obtained using 2d model. Rocking of the fluid was simulated by periodically changed gravity. The model was solved using “Time dependent” solver.

**Discussion S2. Characterization of flow rate on-chip.** The average flow velocity of the device was measured by timing the passage of colored dye.^1, 2^ The flow velocity was measured with five different tilting angles (5⁰, 7.5⁰, 10⁰, 15⁰ and 20⁰) at room temperature. The rocking speed was kept as constant at 0.1 RPM. To carry out the flow visualization, the 3D printed device was placed on the rocker at a neutral position (0⁰ tiling angle). 65 µL of culture medium was slowly added to one side of the medium reservoir using a pipette. Due to the pressure driven by the pipetting and the capillary force within the tubular channel, the medium would flow through the tubular channel and fill both medium reservoirs. 30 µL of the medium was then removed from each reservoir. Due to the surface tension, the tubular channel remained filled with medium. To minimize the disturbance of moving dye front from the pipetting, prior to the loading of colored dye, the tubular device was flash froze with liquid nitrogen for 30 sec. The flash freeze was performed by placing the device on a petri-dish, which was free-floating on liquid nitrogen within a benchtop dwar. Once the device was placed back onto the neutral-positioned rocker. 30 µL of culture medium was quickly added to both sides of the medium reservoir, with one side containing 0.83 mg/mL of Rhodamine B (ChemCruz cat# 203756). The device was then allowed to sit for 1 min at RT to let the frozen medium that was trapped within the tubular channel thaw. One rocking cycle was initiated and the corresponding flow was captured with a fixed positioned video camera on a tripod at 30 frames per second. The flow velocity within the tubular channel was quantified by timing the moving dye front passing through a predetermined distance (3mm). The video was analyzed with ImageJ and the moving dye front was determined with a fixed color intensity threshold. **Fig.S3** shows the representative time-lapse of one rocking cycle at three different tilting angles based on the video recording.

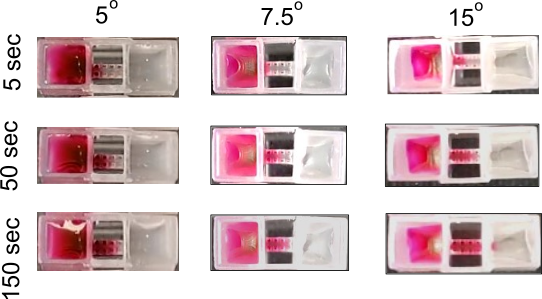

**Fig. S3**: **Experimental characterization of flow rate on-chip.** Representative time-lapse of one rocking cycle of the tubular device at 5⁰, 7.5⁰ and 20⁰, respectively.

**Supplementary Tables**

**Table S1. Medium compositions of cortical organoid protocol**

| **Ingredients** | **Concentrations** |
| --- | --- |
| **Cortical Organoid Medium I (Day 1 – Day 10)** | |
| DMEM/F12 | 1X |
| KOSR | 15% |
| GlutaMax(100X) | 1X |
| MEM-NEAA(100X) | 1X |
| N2 supplement(100X) | 1X |
| b-mercaptoethanol(500X) | 1X |
| Penn/Strep(100X) | 1X |
| SB-431542(10mM) | 1µM |
| Dorsomorphin (10mM) | 2µM |
| XAV939 (30mM) | 2µM |
| **Cortical Organoid Medium II (Day 11 - Day 24)** | |
| DMEM/F12 | 0.5X |
| Neuralbasal | 0.5X |
| N2 supplement(100X) | 1X |
| B27 (wo/VA) (50X) | 1X |
| MEM-NEAA(100X) | 1X |
| GlutaMax(100X) | 1X |
| Insulin (10 mg/mL) | 2.5µg/mL |
| b-mercaptoethanol(500X) | 1X |
| Penn/Strep(100X) | 1X |
| EGF(500µg/mL) | |
| FGF-2(100µg/mL) | 10ng/mL |
| NT3 (100µg/mL) | 10ng/mL |
| BDNF (100µg/mL) | 10ng/mL |
| **Cortical Organoid Medium (Day 25 -)** | |
| DMEM/F12 | 0.5X |
| Neuralbasal | 0.5X |
| N2 supplement(100X) | 1X |
| B27 (w/VA) (50X) | 1X |
| MEM-NEAA(100X) | 1X |
| GlutaMax(100X) | 1X |
| Insulin (10mg/mL) | 2.5µg/mL |
| b-mercaptoethanol(500X) | 1X |
| Penn/Strep(100X) | 1X |
| EGF(500µg/mL) | 10ng/mL |
| FGF-2(100mg/mL) | 10ng/mL |
| NT3 (100µg/mL) | 10ng/mL |
| BDNF (100µg/mL) | 10ng/mL |
| Ascorbic Acid (10 mg/mL) | 0.2mM |
| cAMP (10 mM) | 0.2mM |
| Matrigel | 1% |

**Table S2. Antibody used in immunofluorescence staining**

| **Antigen** | **Host** | **Vendor** | **Catalog#** | **Dilution** |
| --- | --- | --- | --- | --- |
| PAX6 | Rabbit | Biolegend | 901301 | 1:500 |
| MAP2 | Chicken | Millipore | AB5543 | 1:500 |
| CDKN2A/p16INK4a | Mouse | Novus | NBP2-37736 | 1:300 |
| CD11b | Rat | Thermo Fisher | 14-0112-82 | 1:200 |
| GFAP | Rabbit | Abcam | Ab7260 | 1:200 |

**Table S3. Primer sequences for qPCR analysis**

| **Gene** | **Primer sequence** |
| --- | --- |
| hCdkn1a_Fwd | AGG TGG ACC TGG AGA CTC TCA G |
| hCdkn1a_Rev | TCC TCT TGG AGA AGA TCA GCC G |
| hCdkn2a_Fwd | CTC GTG CTG ATG CTA CTG AGG A |
| hCdkn2a_Rev | GGT CGG CGC AGT TGG GCT CC |
| hPTGES2_Fwd | CCA TCC TCT CCC TGG AAA TCT |
| hPTGES2_Rev | ATA TCT GCC GTC CCG AGC TT |
| hPTGER2_Fwd | GAC CAC CTC ATT CTC CTG GCT A |
| hPTGER2_Rev | AAC CTA AGA GCT TGG AGG TCC C |
| hPTGER4_Fwd | AGG CCA TCC GAA TTG CTT CT |
| hPTGER4_Rev | ACT GAG GTC TGG CAG TGA GA |
| hPTGS2_Fwd | CGG TGA AAC TCT GGC TAG ACA G |
| hPTGS2_Rev | GCA AAC CGT AGA TGC TCA GGG A |
| hTNFA_Fwd | CTC TTC TGC CTG CTG CAC TTT G |
| hTNFA_Rev | ATG GGC TAC AGG CTT GTC ACT C |
| hIFNG_Fwd | GAG TGT GGA GAC CAT CAA GGA AG |
| hIFNG_Rev | TGC TTT GCG TTG GAC ATT CAA GTC |
| hIL1B_Fwd | [CAG AAG TAC CTG AGC TCG CC](javascript:openPunchoutItemDetailsForDocLine('PunchoutItemDetailsReqLinePopup',143368255,531204188);) |
| hIL1B_Rev | [AGA TTC GTA GCT GGA TGC CG](javascript:openPunchoutItemDetailsForDocLine('PunchoutItemDetailsReqLinePopup',143368255,531204189);) |
| hARG1_Fwd | [TCA TCT GGG TGG ATG CTC ACA C](javascript:openPunchoutItemDetailsForDocLine('PunchoutItemDetailsReqLinePopup',143368255,531204192);) |
| hARG1_Rev | [GAG AAT CCT GGC ACA TCG GGA A](javascript:openPunchoutItemDetailsForDocLine('PunchoutItemDetailsReqLinePopup',143368255,531204193);) |
| hMCP1_Fwd | [CTC TCG CCT CCA GCA TGA AA](javascript:openPunchoutItemDetailsForDocLine('PunchoutItemDetailsReqLinePopup',143368255,531204194);) |
| hMCP1_Rev | [GAA GAA GAG GGG GCC TTA CC](javascript:openPunchoutItemDetailsForDocLine('PunchoutItemDetailsReqLinePopup',143368255,531204195);) |
| hLIF_Fwd | [AGA TCA GGA GCC AAC TGG CAC A](javascript:openPunchoutItemDetailsForDocLine('PunchoutItemDetailsReqLinePopup',143368255,531204196);) |
| hLIF_Rev | [GCC ACA TAG CTT GTC CAG GTT G](javascript:openPunchoutItemDetailsForDocLine('PunchoutItemDetailsReqLinePopup',143368255,531204197);) |
| hMCP3_Fwd | [ACA GAA GGA CCA CCA GTA GCC A](javascript:openPunchoutItemDetailsForDocLine('PunchoutItemDetailsReqLinePopup',143368255,531204198);) |
| hMCP3_Rev | GGT GCT TCA TAA AGT CCT GGA CC |
| hMCP2_Fwd | [TAT CCA GAG GCT GGA GAG CTA C](javascript:openPunchoutItemDetailsForDocLine('PunchoutItemDetailsReqLinePopup',143368255,531204200);) |
| hMCP2_Rev | [TGG AAT CCC TGA CCC ATC TCT C](javascript:openPunchoutItemDetailsForDocLine('PunchoutItemDetailsReqLinePopup',143368255,531204201);) |
| hRANTES_Fwd | [TCA TTG CTA CTG CCC](javascript:openPunchoutItemDetailsForDocLine('PunchoutItemDetailsReqLinePopup',143368255,531204202);) |
| hRANTES_Rev | [TCG GGT GAC AAA GAC](javascript:openPunchoutItemDetailsForDocLine('PunchoutItemDetailsReqLinePopup',143368255,531204203);) |
| hTOMM20_Fwd | [CGA CCG CAA AAG ACG](javascript:openPunchoutItemDetailsForDocLine('PunchoutItemDetailsReqLinePopup',143368255,531204204);) |
| hTOMM20_Rev | [GCT TCA GCA TCT TTA](javascript:openPunchoutItemDetailsForDocLine('PunchoutItemDetailsReqLinePopup',143368255,531204205);) |
| hVDAC1_Fwd | [GCA AAA TCC CGA GTG](javascript:openPunchoutItemDetailsForDocLine('PunchoutItemDetailsReqLinePopup',143368255,531204206);) |
| hVDAC1_Rev | [TCC AGG CAA GAT TGA CAG CGG T](javascript:openPunchoutItemDetailsForDocLine('PunchoutItemDetailsReqLinePopup',143368255,531204207);) |
| hSLC13a3_Fwd | [CCA TTG AGG AGT GGA](javascript:openPunchoutItemDetailsForDocLine('PunchoutItemDetailsReqLinePopup',143368255,531204208);) |
| hSLC13a3_Rev | [GTG TTG CTC AGC CAC](javascript:openPunchoutItemDetailsForDocLine('PunchoutItemDetailsReqLinePopup',143368255,531204209);) |
| hNOS2_Fwd | [GCT CTA CAC CTC CAA TGT GAC C](javascript:openPunchoutItemDetailsForDocLine('PunchoutItemDetailsReqLinePopup',143368255,531204210);) |
| hNOS2_Rev | [CTG CCG AGA TTT GAG CCT CAT G](javascript:openPunchoutItemDetailsForDocLine('PunchoutItemDetailsReqLinePopup',143368255,531204211);) |
| hGFAP_Fwd | [CTG GAG AGG AAG ATT GAG TCG C](javascript:openPunchoutItemDetailsForDocLine('PunchoutItemDetailsReqLinePopup',143368255,531204212);) |
| hGFAP_Rev | [ACG TCA AGC TCC ACA TGG ACC T](javascript:openPunchoutItemDetailsForDocLine('PunchoutItemDetailsReqLinePopup',143368255,531204213);) |
| hSERPINA3_Fwd | [CCT GAA CGA CAT ACT](javascript:openPunchoutItemDetailsForDocLine('PunchoutItemDetailsReqLinePopup',143368255,531204214);) |
| hSERPINA3_Rev | [CAT CAA GCA CAG CCT](javascript:openPunchoutItemDetailsForDocLine('PunchoutItemDetailsReqLinePopup',143368255,531204215);) |
| hVimentin_Fwd | [AGG CAA AGC AGG AGT](javascript:openPunchoutItemDetailsForDocLine('PunchoutItemDetailsReqLinePopup',143368255,531204216);) |
| hVimentin_Rev | [ATC TGG CGT TCC AGG](javascript:openPunchoutItemDetailsForDocLine('PunchoutItemDetailsReqLinePopup',143368255,531204217);) |
| hNestin_Fwd | [TCA AGA TGT CCC TCA GCC TGG A](javascript:openPunchoutItemDetailsForDocLine('PunchoutItemDetailsReqLinePopup',143368255,531204218);) |
| hNestin_Rev | [AAG CTG AGG GAA GTC TTG GAG C](javascript:openPunchoutItemDetailsForDocLine('PunchoutItemDetailsReqLinePopup',143368255,531204219);) |
| GAPDH_Fwd | GAC AGT CAG CCG CAT CTT CT |
| GAPDH_Rev | AAA TGA GCC CCA GCC TTC TC |

**References**

1.	Wang, Y.I. & Shuler, M.L. UniChip enables long-term recirculating unidirectional perfusion with gravity-driven flow for microphysiological systems. *Lab Chip* **18**, 2563-2574 (2018).

2.	Lin, T.Y., Do, T., Kwon, P. & Lillehoj, P.B. 3D printed metal molds for hot embossing plastic microfluidic devices. *Lab Chip* **17**, 241-247 (2017).
